# Doxycycline Modulates Uveal-Melanoma-Associated Marker Expression in BAP1-Repressed Human Ocular Organoids

**DOI:** 10.64898/2026.07.26.740828

**Authors:** Ethan A Chiu, Timothy A Blenkinsop

**Affiliations:** Department of Ophthalmology, Cell Development and Regenerative Biology, Black Family Stem Cell Institute, Icahn School of Medicine at Mount Sinai, New York, NY 10029

**Keywords:** Uveal Melanoma, Doxycycline, RNA Sequencing, SEAM Organoid, Immunofluorescence, RT-qPCR, CRISPR-Cas9, organoid, human stem cell, doxycycline, neural crest

## Abstract

Uveal Melanoma (UM) is the most common eye cancer, with a metastatic mortality rate of 80%. Only 1-3% of patients have detectable UM at metastasis, and UM exhibits punctuated early growth. Doxycycline has recently been shown to inhibit metabolic processes exploited by cancer cells and reduce cancer cell growth in models of liver cancer. We hypothesized doxycycline may also be effective in UM and therefore tested doxycycline treatment in an eye organoid model of uveal melanoma. Using a stem cell line whereby BAP1 can be knocked down with a tetracycline-inducible system, we differentiated this line into a whole eye organoid model termed self-formed ectodermal autonomous multi-zone of ocular cells (SEAM). We found an enhanced proliferation in neural crest cells within the SEAM colonies. To identify the neural crest cells, we conducted single-cell RNA sequencing (scRNA-seq) analysis utilizing the Seurat R toolkit to pinpoint genes within neural crest clusters. To confirm the results of the *in silico* scRNA-seq analysis, genes with notable functions and differential expression in the neural crest cluster in relation to UM proliferation, angiogenesis, and oxidative phosphorylation were analyzed through immunofluorescence and RT-qPCR. Based on the scRNA-seq analysis, immunofluorescence, and RT-qPCR, the novel BAP1 KD (UM phenotype) model was found to replicate UM-relevant gene and protein expressions effectively, so the BAP1 KD (UM phenotype) was then treated with doxycycline to evaluate its effect on UM metastasis. Subsequent analysis found that doxycycline significantly inhibited UM growth, angiogenesis, and oxidative phosphorylation in the BAP1 KD (UM phenotype) model more than that of the control model, perhaps due to doxycycline targeting higher regions with more mitochondrial activity, indicating doxycycline’s therapeutic potential in treating UM.

## Introduction

Uveal melanoma (UM) is the most common cancer of the eye and the second most common form of melanoma. The four-year mortality rate of patients with metastatic UM is approximately 80% (Rietschel, 2005), with a median survival period of fewer than six months (Singh, 2005). Only 1-3% of patients have detectable metastases at diagnosis (Rietschel, 2005), and UM exhibits punctuated early growth (Field, 2018). Thus, rapid and early treatment is critical for long-term survival. Unfortunately, within 2.4 years of primary tumor treatment, 50% of UM patients develop metastasis (Nabil, 2015), with 95% of metastases to the liver (Bellerive, 2018). Therefore, the minority of UM patients successfully treated after their initial occurrence of UM are still at long-term risk (Kolandjian, 2013). Furthermore, current therapies like chemotherapy and enucleation are ineffective. Chemotherapy has negligible impacts on overall UM patient survival (Afzal, 2018), with only a 0-6% success rate (Kinsey, 2017), and leads to adverse side effects including vomiting and nausea in 40-50% of patients (Rodriguez-Vidal, 2020). Enucleation removes the entire eye, resulting in vision loss, and has a 5-year mortality rate of 50% (Krohn, 2007). Given the poor prognosis of those diagnosed with UM and the lack of a documented cure, understanding the origin and mechanism behind UM cell proliferation is essential to developing effective treatments. Specifically, BAP1, SF3B1, or EIF1AX gene mutations are critical to UM proliferation (Field, 2018), with BAP1 more prevalent in the more deadly UM class 2 variant, and SF3B1 and EIF1AZ mutations more prevalent in the UM class 1 variant. BAP1 loss also leads to tumor cells developing the stem cell-like properties associated with the neural crest (Kuznetsov, 2019).

While animal and human cell lines are most frequently used to test UM treatments, animals have different genetic makeups than humans, leading to potential inaccuracies. Furthermore, human tissue samples cannot be controlled for and are not only highly individual but also prone to contamination and mutation, introducing potential bias and preventing universally reproducible results. Newly developed human stem cell-derived models more closely resemble human cell physiology, and organoid models have been found to resemble human organs. Thus, they can be used as avatars or sentinels for patients with diseases. One such model, the self-formed ectodermal autonomous multi-zone (SEAM) organoid developed by Hayashi (2016), is an *in vitro* model of eye development with the potential to overcome the limitations of animal experiments and human cell lines (Eriksen, 2021).

Single-cell transcriptional analysis (scRNA-seq) reveals the gene expression profiles of individual cells and is currently the best method to define cell states and phenotypes (Tanay, 2017). Currently, Seurat is the most accurate and efficient scRNA-seq analysis toolkit (Zhao, 2019). Since gene expression patterns can be identified using cell clustering analysis, differentially expressed genes between the SEAM organoid and UM cells of varying types can be mapped, facilitating the location of diagnostic and therapeutic targets for various eye-related diseases.

Interestingly, Matsumoto (2017) found that the prescription antibiotic doxycycline decreased oxidative phosphorylation through mitochondrial biogenesis inhibition, limiting mitochondria-dependent cancer stem cell proliferation. Moreover, Artegiani (2019) noted that doxycycline acutely increased tumor suppressors such as BAP1 that control metastasis in human liver organoids, where UM often metastasizes. Considering these studies and how expensive and ineffective current therapies are for UM, the generic antibiotic doxycycline is an effective, efficient, and economical treatment candidate. The current study analyzed doxycycline’s effects on markers affecting tumor suppression, tumor growth, mitochondrial oxidative phosphorylation, and angiogenesis using immunofluorescence, which quantifies protein expression, and quantitative reverse transcription polymerase chain reaction (RT-qPCR), which quantifies gene expression.

To introduce targeted gene knockdowns (KDs) into organoids and cell culture tissues, lentiviral vectors can be used to deliver the CRISPR guide RNAs that aid Cas9 nucleases in DNA cleavage (Milone, 2018). Lentiviral transduction was used to introduce gene KDs, such as those targeting tumor suppressor genes (Moses, 2020). Thus, cell proliferation indicative of tumor growth can be induced in the SEAM organoid by inhibiting crucial cell regulator genes. Ultimately, the UM replication potential of a model created through the KD of the commonly mutated UM tumor suppressor gene BAP1 was evaluated, and doxycycline’s effects on the UM metastasis phenotype organoids were compared to those of the control and doxycycline-free BAP1 KD organoid using immunofluorescence and RT-qPCR.

## Methodology

### Seurat scRNA-seq analysish

Following RNA isolation of single cells from an H9 SEAM ocular eye model organoid and the creation of an RNA sequencing library for the single cells, a sequenced SEAM dataset was created using the Illumina NovaSeq 6000 Sequencing System and mapped to a human reference genome (Haque, 2017). The dplyr, Seurat (Hao, 2021), and patchwork libraries, along with the SEAM dataset, normal human eye data, and UM cell models (Harbour, 2020), were loaded into RStudio (RStudio Team, 2021) with R version 4.1.0 (R Core Team, 2021). The resulting Seurat object was initialized with the raw data. By visualizing the quality control metrics using a violin and feature scatter plot to exclude empty droplets with very few genes, cell multiplets with an abnormally high gene count and dying cells featuring burst mitochondria were filtered out of each investigated dataset. The feature expression measurements were normalized using the total expression, and the result was log-transformed. To shift gene expression so highly expressed genes would not dominate by making the mean-variance zero across all cells and the cell variance one, linear transformation scaling was applied. Next, principal component analysis linear dimensional reduction was performed to increase interpretability and minimize information loss by creating uncorrelated variables to maximize variance. Significant PCs with low p-value features were identified by observing the ElbowPlot function to find the true dimensionality of the dataset. The FindNeighbors function was run to cluster the genes by finding the distance between neighboring genes. Furthermore, the FindCluster function was run to group genes together using the Louvain algorithm. To create a dimensional plot using UMAP reduction, non-linear dimensionality reduction was run using the RunUMAP function. Lastly, the positive markers for every cluster with a minimum percentage of 0.25 and a log fold change threshold of 0.25 were found, and the resulting clusters were grouped together. These clusters were visualized through a UMAP plot.

### Cluster identification

The seam.markers dataset created from the scRNA-seq analysis was exported to a .csv file, and all the genes from each cluster within the .csv file were copied onto a spreadsheet. Following the separation of each cluster onto a different tab, the genes from each cluster were copied and inputted into Enrichr (Xie, 2021) to identify the probable cluster identities. Screenshots of the Mouse Gene Atlas Table and Human Gene Atlas Table were inserted into their respective cluster tabs. Subsequently, a literature review was conducted and the GeneCards (Stelzer, 2016) human genome database was consulted to find specific canonical gene markers identifying each region of the eye. The gene markers identified were gathered into separate spreadsheet tabs. Identified gene markers were used to classify clusters as specific regions of the eye.

### Seurat scRNA-seq cluster labeling

The cell-type identity was assigned to cell clusters using the new.clusters.ids function, and an annotated UMAP plot was created. The accuracy of the SEAM model in representing the human eye was determined based on cluster identities and comparison with an actual sequenced human eye sample. Based on the presence of neural crest cluster identities, the potential of using the SEAM model to investigate UM was evaluated. The location and number of notable differentially expressed UM genes were noted in the normal human eye cells, the UM tumor cells, and SEAM eye organoid clusters with an emphasis on the neural crest clusters since the neural crest is a common location from which UM gene mutations leading to metastasis may arise.

### Maintaining Cell Cultures

dCas9 KRAB hiPSC stem cells were obtained from the Eye Bank for Sight Restoration (AICS-0090 cl.391). The stem cells were cultured and maintained within the 6-well plates (Corning) in a cell culture hood (The Baker Company). Every two days, 2mL of mTeSR Plus complete media, made up of 40mL mTeSR Plus (Stemcell Technologies), 10mL mTeSR Plus 5X supplement (Stemcell Technologies), 0.5mL glutamax (Gibco), and 0.5mL penicillin-streptomycin (Gibco), was changed. Typically, the cells became 70-80% confluent after five days in mTeSR Plus complete media. To prevent over-confluence and differentiation, the stem cells were passaged and subcultured every five days.

Wells within a 6-well plate were coated with 0.15mL Matrigel (Corning) dissolved in 11.85mL 1:1 Dulbecco’s Modified Eagle Medium/Nutrient Mixture F-12 (Corning) media. Well plates needed immediately were placed in the incubator (Thermo Scientific) at 37.0°C, 5.0% carbon dioxide, and 21.0% oxygen for 1 hour. Otherwise, well plates were stored at 3.8°C. The calcium and magnesium bonds between the stem cells and the well plate were chelated to harvest the maintenance cells. Next, the maintenance culture was incubated at 37.0°C, 5.0% carbon dioxide, and 21.0% oxygen in 1mL of 0.5mM ethylenediaminetetraacetic acid (EDTA) (Corning) for 5 minutes. The EDTA was aspirated, and then a 1mL of the mTeSR Plus complete and a 0.2μL dilution of thiazovivin (Stemcell Technologies) Rho-associated protein kinase inhibitor solution was placed into the maintenance well. The plate was agitated to lift cells into the media, and the resulting media was split 1:4 into wells in a new 6-well plate. The new plate was placed in the incubator (Thermo Scientific) and left overnight for cell-plate adhesion.

### Creating SEAM organoids

To create SEAM organoids from cultured stem cells (Eriksen, 2022), the subculturing steps for maintaining the stem cells were followed with the exception of the 1:4 split. Instead, to determine the number of cells in each well and the subsequent split ratio, 10µl of trypan blue (Gibco) and 10µl of maintenance cell solution were pipetted onto the hemocytometer (Hausser Scientific). The inverted microscope (Leica) at 5x was used to count cells in the central and four corner squares, and the following formulas were used to calculate the total cells per mL:

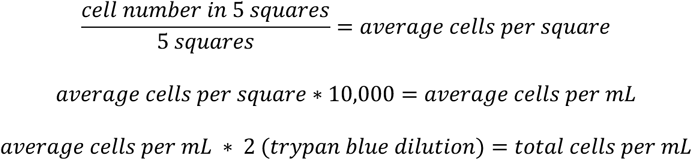

Since the optimal SEAM organoid split is 2,500 cells per mL (Eriksen, 2021) and all wells in the 6-well plate were used to grow SEAM organoids, the ideal number of cells per mL was 15,000.

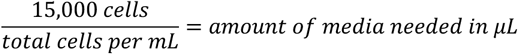

The calculated media volume was divided among each well in the 6-well plate, placed in the incubator at 37.0°C, 5.0% carbon dioxide, and 21.0% oxygen, and left overnight for cell-plate adhesion.

SEAM media was made using 225mL Glasgow’s Minimal Essential Medium (GMEM) (Gibco), 25mL KnockOut Serum Replacement (KSR) (Life Technologies), 2.5mL sodium pyruvate (Gibco), 2.5mL minimum essential medium non-essential amino acids (Gibco), 2.5mL glutamax, 2.5mL penicillin-streptomycin, and 96µL 2-Mercaptoethanol (Sigma-Aldrich). After waiting 4-5 days for the stem cells to reach 70-80% confluence, feeding the stem cells mTeSR Plus complete every two days in between, the stem cells were slowly transitioned to SEAM media by feeding them with 1mL of SEAM media and 1mL of mTeSR Plus complete media for two cycles before converting to SEAM media only. The development of the SEAM cultures was periodically monitored through the inverted microscope at 5x. At 35 days old, the SEAM cultures were considered mature and ready for analysis.

### Lentiviral transduction

To reduce cell adhesion, dCas9 KRAB cells were given 2mL (6-well plate) of Dulbecco’s phosphate-buffered saline (PBS), which is calcium-magnesium free. For each BAP1 KD lentiviral condition — green fluorescent protein (GFP) control and GFP + doxycycline — 0.5µL/mL of puromycin was added to the media in preparation for selecting cells that are stably expressing shRNA for the BAP1 KD. To minimize viral and cell surface repulsion (Moses, 2020) and to improve transduction efficiency (Horani, 2013) respectively, 8mg/μL of polybrene (Sigma Aldrich) and a 1:5000 dilution of thiazovivin were added to the combined solution. For each dCas9 KRAB SEAM well, 30µL of lentivirus (Feng Zhang Lab) was added. The 6-plate was parafilmed and placed into a centrifuge (Beckman Coulter) at 21°C room temperature and 1000 rpm (205 × g) for 90 minutes to concentrate the lentiviral particles near the SEAM organoids (Horani, 2013). Next, the organoids were placed into the incubator for 48 hours.

### Treating SEAM organoids with doxycycline

After 36 hours of incubation, a 1:1000 dilution of doxycycline to SEAM media was inserted into the non-lentiviral-infected doxycycline control and BAP1 KD GFP + doxycycline wells. Basic SEAM solution was fed to the non-lentiviral-infected control and BAP1 KD GDP control. To further aid in lentiviral penetration, 0.5µL/mL of puromycin was added to all the media. After 24 hours of doxycycline incubation, the doxycycline-treated cells were switched to puromycin-free doxycycline-infused SEAM media. After these steps, the doxycycline-treated cells were then provided doxycycline-infused SEAM media every 48 hours for the next 14 days following the lentiviral transduction.

### Fixing SEAM samples

After 11 days — totaling 14 days since the lentiviral transduction — all SEAM cell culture wells were rinsed three times with 2mL of PBS (Corning) with 5 minutes between each rinse. Next, the SEAM cell cultures were treated with paraformaldehyde (Fisher Scientific) for 20 minutes. The SEAM cell cultures were then rinsed thrice with PBS in the chemical hood with 5 minutes in between to remove the paraformaldehyde.

### Permeabilization and blocking

A solution of 0.2% Triton-X-100 (Fisher Bioreagents), which punctures the cell membranes to allow primary antibodies to enter, 5% normal donkey serum (Fisher Scientific), which blocks non-specific antibody binding, 1% bovine serum albumin (BSA) (Fisher Bioreagents), which reduces imaging background, and PBS was created in a 15mL tube (Corning). Each well was given 2mL of solution and left to permeabilize and block for 30 minutes.

### Primary antibody staining

To remove the permeabilization and blocking solution, the SEAM cell cultures were rinsed three times with 2mL of 1% BSA in PBS with 5 minutes between each rinse. Four different SEAM well conditions were tested. The primary antibodies were diluted with 2mL of 1% BSA in PBS with 1mL for each combination, allowing a quarter to be given to the non-doxycycline-treated negative control, doxycycline control, BAP1 KD GFP control, and BAP1 KD GFP + doxycycline wells. For each control and experimental group, 4μL of STRO1 (CD34) (mouse, Abcam, ab102969, 1:250) antibody, 40μL of KI67 (rabbit, Abcam, ab16667, 1:25) antibody, and 10μL of ABCB5 (goat, Abcam, Ab77549, 1:100) antibody were placed in well 1, given based on the manufacturer’s suggested dilution. For well 2, 2μL of BAP1 (mouse, Abcam, ab167250, 1:500) antibody, 40μL of KI67 (rabbit, Abcam, ab16667, 1:25) antibody, and 10μL of ABCB5 (goat, Abcam, Ab77549, 1:100) antibody were similarly provided. Additionally, 3.3μL of PAX3 (mouse, DSHB, clone unspecified, lab aliquot) antibody, 20μL of AKT1 (rabbit, Abcam, ab235958, 1:50) antibody, and 10μL of ABCB5 (goat, Abcam, Ab77549, 1:100) antibody were provided for well 3. For well 4, 10μL of ABCB5 (goat, Abcam, Ab77549, 1:100) antibody, 2μL of SOX10 (mouse, R&D Systems, MAB2864, 1:500) antibody, and 10μL of CASP3 (rabbit, Thermo Fisher, 700182, 1:100) antibody were given. After adding the corresponding antibodies to each well, the 6-well plates were wrapped with parafilm and left on the 3D Platform Rotator (Fisherbrand) at 3.8°C overnight.

### Secondary antibody staining

The SEAM cultures were rinsed with 2mL of 1% BSA in PBS three times with 5 minutes in between each rinse to remove the primary antibody solutions. Two different conditions were created for the secondary antibody staining. Since all wells use rabbit, mouse, and goat antibodies, 0.5μL of DAPI (Thermo Scientific, 62248, 1:200), 0.5μL of rabbit Alexa Fluor 546 (Invitrogen, A11071, 1:200), 0.5μL of goat Alexa Fluor 647 (Invitrogen, A21447, 1:200), and 0.5μL of mouse Alexa Fluor 488 (Invitrogen, A11001, 1:200) were added to 2mL of 1% BSA in PBS, with 100μL going to each well. After being wrapped in parafilm and stored in darkness for one hour at 3.8°C, each SEAM well was washed with 100μL of 1% BSA in PBS three times with 5 minutes between each rinse to remove the secondary antibody solutions. After the third rinse, 100μL of 1% BSA in PBS was maintained in each well to prevent the SEAM organoids from drying out. If not needed immediately, the well plates were parafilmed and stored at 3.8°C.

### Immunofluorescence microscopy

An immunofluorescence microscope (Leica) featuring a motorized stage control and DAPI, GFP 488, RFP/Cy3 546, Cy5 647, and phase filters controlled by Leica Application Suite X were used to analyze the protein expression. Several micrometers of the SEAM organoid were scanned to capture the images. The exposure time for the images was determined through trial and error and standardized across all the wells, with the maximum pixel intensity aligned with the fluorescence reference intensity’s linear range (Kamao, 2014). Immunoreactive cells were uniformly counted in randomly chosen areas (6-well plate) using ImageJ (Grishagin, 2015). Statistical comparisons between the two groups were made using an Independent Samples t-test.

### Immunofluorescence analysis

After the protein expression images were derived from immunofluorescence microscopy, the cells indicated by blue dots or outlines were counted through ImageJ (Labno, 2020). For all protein markers except for ABCB5, which was indicated by colored cell outlines, protein expression was identified by counting the dots identifying nuclei. For the GFP groups, more pronounced dots indicated lentiviral infection in the nucleus. To count the cells expressing the specific protein markers, the protein-specific images were converted to 16-bit greyscale. Next, the image threshold was adjusted to remove the background and highlight the cells to be counted. If the resulting particles were touching, the background was subtracted to reduce noise and background pixels. Thus, a binary image was created with only black-and-white pixel intensities. If the particles merged following the binary image creation, watershed separation was run to separate the particles. Finally, the ‘analyze particles’ function was run with the size set between 110 and 250 square pixels to account for small and abnormal metastasized cell nuclei. The circularity was adjusted to 0.50-1.00 to reduce error and ensure that the cells detected were of a shape between an imperfect circle and a perfect circle instead of a straight line. To identify KI67 + STRO1 positive co-expression, the KI67 and STRO1 images were overlaid. The protein fluorescence intensity for each image was also measured in ImageJ to corroborate the cell counting results.

For each protein marker, cell counts and mean fluorescence intensity were quantified from the immunofluorescence images. Measurements from multiple images obtained within the same experimental replicate were first averaged to generate a single value for that replicate. The mean and standard deviation were then calculated across independent experimental replicates. The standard error of the mean (SEM) was calculated as the standard deviation divided by the square root of the number of independent experimental replicates. Bar charts were generated to display the mean cell count or fluorescence intensity for each condition, with error bars representing the SEM.

### RNA isolation

The SEAM media was removed from four wells for each condition and rinsed with 2mL each of CMF PBS for 30 seconds. A mixture of 1mg/mL of DNAse and 2mL of CMF PBS was added to each well, and each well was incubated at 37°C for 3 minutes. The SEAM cultures were scraped off the plates and placed into 15mL tubes. Subsequently, the 15mL tubes were placed in a centrifuge (Beckman Coulter) for 5 minutes at 5000 rpm (3070 × g). After removing the supernatant, the pellets were incubated in 1mL of RNAprotect (Qiagen) at 37°C for 5 minutes. To optimize RNAprotect penetration, the contents were transferred to 1.5mL Eppendorf tubes, titrated, and spun in a centrifuge (Eppendorf MiniSpin plus) for 10 minutes at 14,500 rpm (14100 × g). Extra RNA was stored at −80°C. The Qiagen RNeasy Micro Kit (Cat. 74004) was used to extract the RNA. The four resulting RNA tubes were incubated at 37°C for 10 minutes to increase RNA yield. The resulting concentrations of the purified RNA samples were quantified using a Thermo Scientific NanoDrop ND-1000 spectrophotometer. RNA concentration values for each sample can be found in Supplemental Table 1.

### Complementary DNA (cDNA) conversion

The Applied Biosystems High-Capacity cDNA Reverse Transcription Kit (Cat. 4368814) was used to create the master mix for cDNA conversion. For each tube, 6μL of master mix and 14μL of RNA sample were combined. The combined solution was placed in a thermocycler (Eppendorf) at 37°C for 60 minutes, 95°C for 5 minutes, and 3.8°C for 30 minutes to activate the reverse transcriptase enzyme. After cDNA conversion, molecular grade water was added to ensure the resulting cDNA in each of the four samples was converted to 10ng/mL based on the RNA concentration values (assuming a 1:1 input-to-output ratio).

### RT-qPCR gene detection and quantification

Forward and reverse cyclophilin A, PRKDC, BAP1, p53, ABCB5, PAX3, PGC1α, NRF1, HIF1, and SIRT1 (Integrated DNA Technologies) primers were placed in a 1:100 dilution in an Eppendorf tube with 98μL molecular grade water and 1μL of both forward and reverse primer. The tube contents were placed into a master mix tube (Eppendorf) with SYBR Green (Alkali Scientific) in a 1:10 dilution. The resulting combined primer and master mix solution was sectionally placed onto the 384 well plate (Applied Biosystems) at 3μL for each well. The cDNA samples were placed into the top rows of the 384 wells at 1μL for each well. To account for potential errors, every gene was allocated three wells. A MicroAmp Optical Adhesive Film (Applied Biosystems) was used to seal the 384 well plate. RT-qPCR was run on an Applied Biosystems QuantStudio 6 Pro using primers. RNA levels were normalized to the cyclophilin A housekeeping gene. Next, cycles were run for each gene and sample group until the fluorescent signal crossed the threshold intensity, which was then recorded as the quantification cycle (Cq) values.

### RT-qPCR analysis

Following the export of the raw RT-qPCR data from the Applied Biosystems QuantStudio 6 Pro, the sample, target, and quantification cycle (Cq) values were evaluated for each gene and sample group. For each sample, the average Cq value of the three wells featuring each gene was calculated as the mean Cq. Additionally, the average Cq indicating cyclophilin gene expression for each sample was determined as the cyclophilin mean Cq. To calculate the change in cycle threshold (delta Ct), which is equivalent to the change in Cq (delta Cq), the sample cyclophilin mean Cq was subtracted from the evaluated gene mean Cq. To find the delta-delta Ct values, which determine the relative fold gene expression for each sample, the calculated delta Ct values for the control were subtracted from the calculated delta Ct values of each experimental group. Next, the relative quantification was determined by raising 2 to the power of the delta-delta Ct value (2^−ΔΔCt^). The resulting relative quantification was used to determine significant changes in gene expression levels between each experimental group and the corresponding sample control.

For each biological replicate, the Cq values from the three technical wells were averaged separately for each target gene and for the cyclophilin A reference gene. The ΔCq value was calculated by subtracting the mean cyclophilin A Cq from the mean target-gene Cq. Within each experiment, the ΔΔCq value was calculated by subtracting the ΔCq of the matched control condition from the ΔCq of each experimental condition. Relative gene expression was then calculated using the 2^-ΔΔCq method. Relative-expression values were summarized across the five independent experimental replicates. Bar charts display the mean relative expression for each condition, with error bars representing the SEM, calculated as the standard deviation divided by the square root of the number of independent experimental replicates. The three technical wells were averaged and were not treated as independent observations.

## Results

The SEAM scRNAseq raw data was sequenced in the Blenkinsop lab. The UM class 1 metastatic, UM class 2 primary, and UM class 2 metastatic scRNA-seq data were derived from that published by Durante 2020. The original raw data was downloaded from the Gene Expression Omnibus (dbGaP accession phs001861.v1.p1) and loaded using Seurat.

dCas9-KRAB stem cell cultures were differentiated into SEAM eye organoids. Next, scRNA-seq facilitated SEAM transcriptome mapping to a human reference genome. To increase interpretability and minimize information loss, principal component analysis (PCA) of single-cell transcript reads was performed through RStudio. Following normalization and quality control, cell clusters were created by grouping SEAM cells of similar gene expressions.

The UM class 1 metastatic (BSSR0022) had one raw dataset. The multiple UM class 2 primary (UMM059, UMM061, UMM062, UMM063, UMM064, UMM065, UMM066, UMM069) and UM class 2 metastatic (UMM041L, UMM067L) raw datasets were optimally combined for the most accurate reading of differentially expressed genes. PCA of single-cell transcript reads, normalization, quality control, and cell clustering were similarly performed on all three datasets.

The SEAM, UM class 1 primary, UM class 1 metastasis, and UM class 2 metastasis clusters were organized into a spreadsheet (Figure 1) and identified based on their region of origin in the eye using gene markers from literature reviews, the Enrichr gene list analysis tool (Figure 2), and the GeneCards database. Subsequently, the clusters were labeled onto UMAP plots using the scRNA-seq analysis program (Figure 3). Significantly differing (p<0.05) gene expressions were found among corresponding normal human eye, SEAM, and UM clusters of various classes using the Seurat Wilcoxon rank sum test, and likely genes involved in metastasis were labeled and identified within the UM clusters.

**Figure 1:**
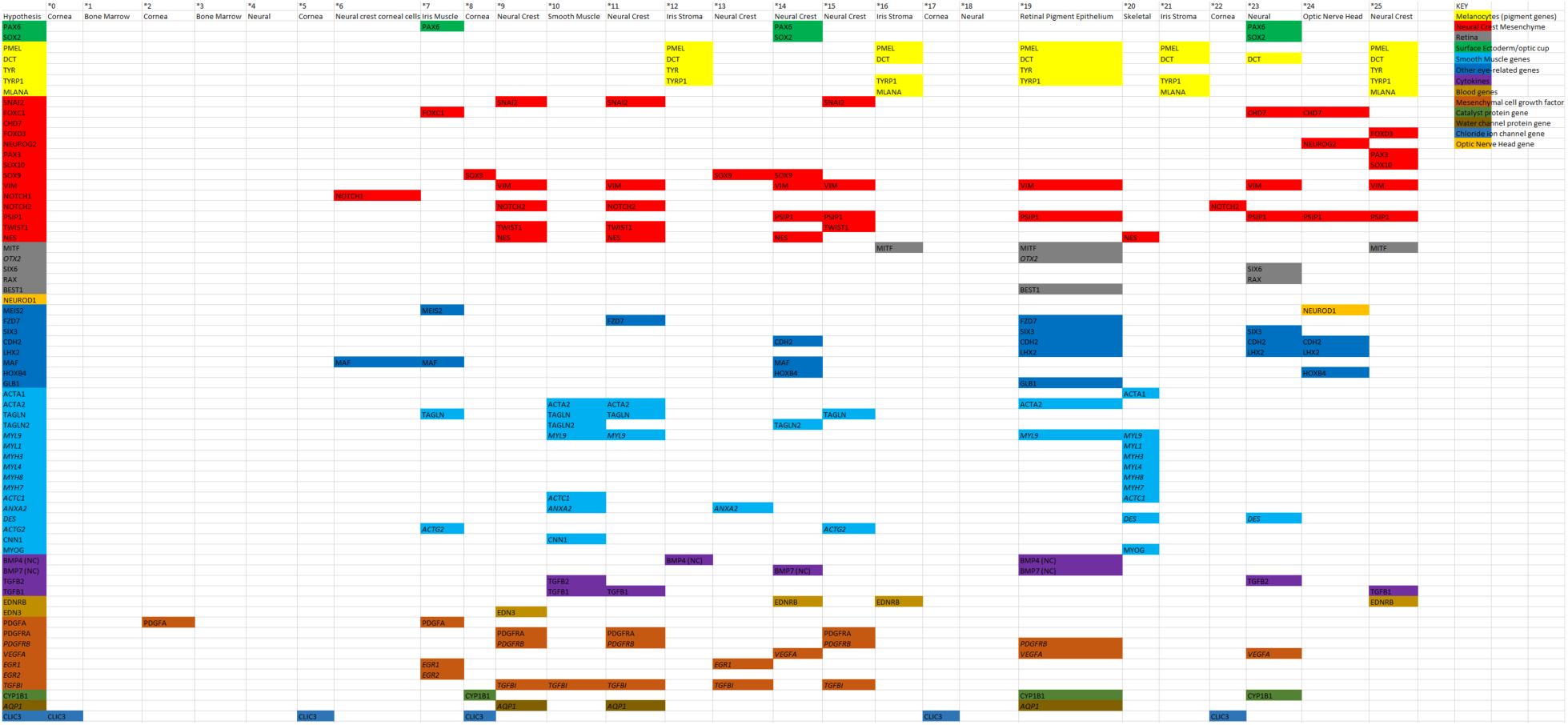
Excel spreadsheet documenting gene function groupings and labeling of the scRNA-seq-derived cell clusters. The clusters were identified using canonical gene markers, gene function literature reviews, Enrichr, and GeneCards analysis. Genes (rows) with a similar function were color-coded, and various functions combined allowed for the clusters (columns) to be labeled correspondingly.

**Figure 2:**
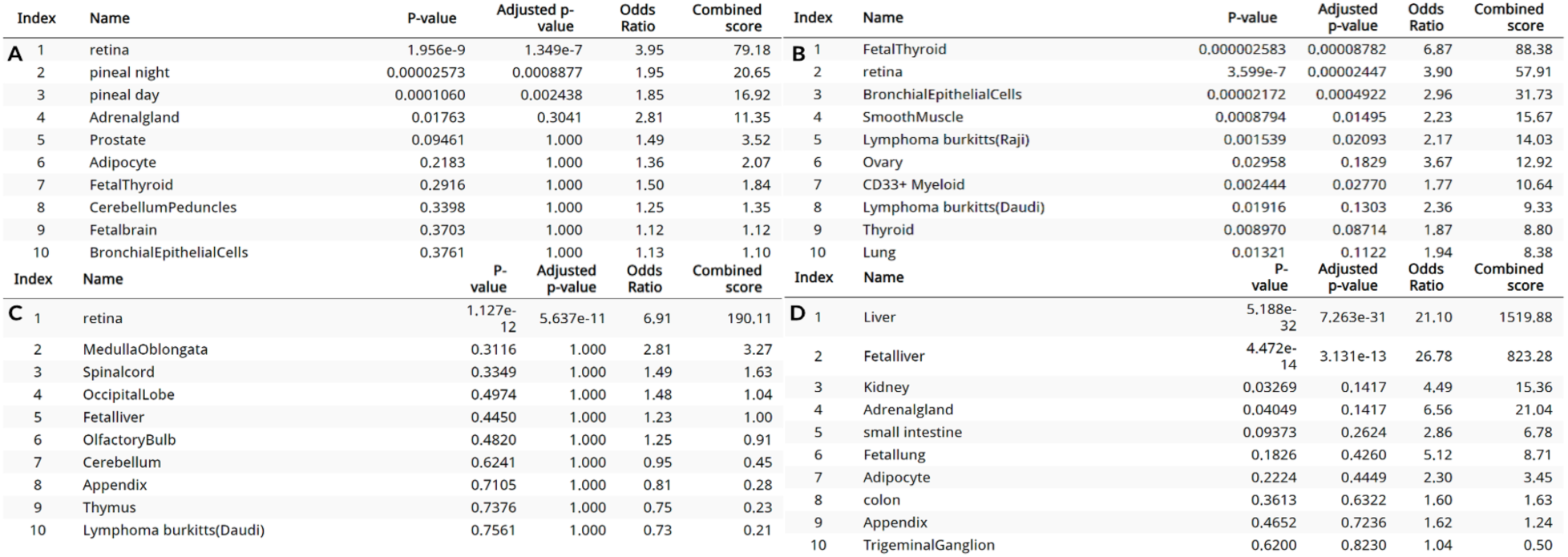
Statistically significant Enrichr human gene atlas predictions that aid in determining cluster identities for the normal human eye, SEAM organoid, and UM phenotypes. Enrichr human gene atlas predictions for specific clusters within the (A) SEAM organoid, (B) UM class 1 metastatic, (C) UM class 2 primary, and (D) UM class 2 metastatic phenotypes. Notably, Enrichr prediction of the UM class 2 metastasis phenotype reveals that the genes in the cluster are indicative of the liver rather than the retinas predicted in the SEAM, UM class 1 metastasis, and UM class 2 primary models.

**Figure 3:**
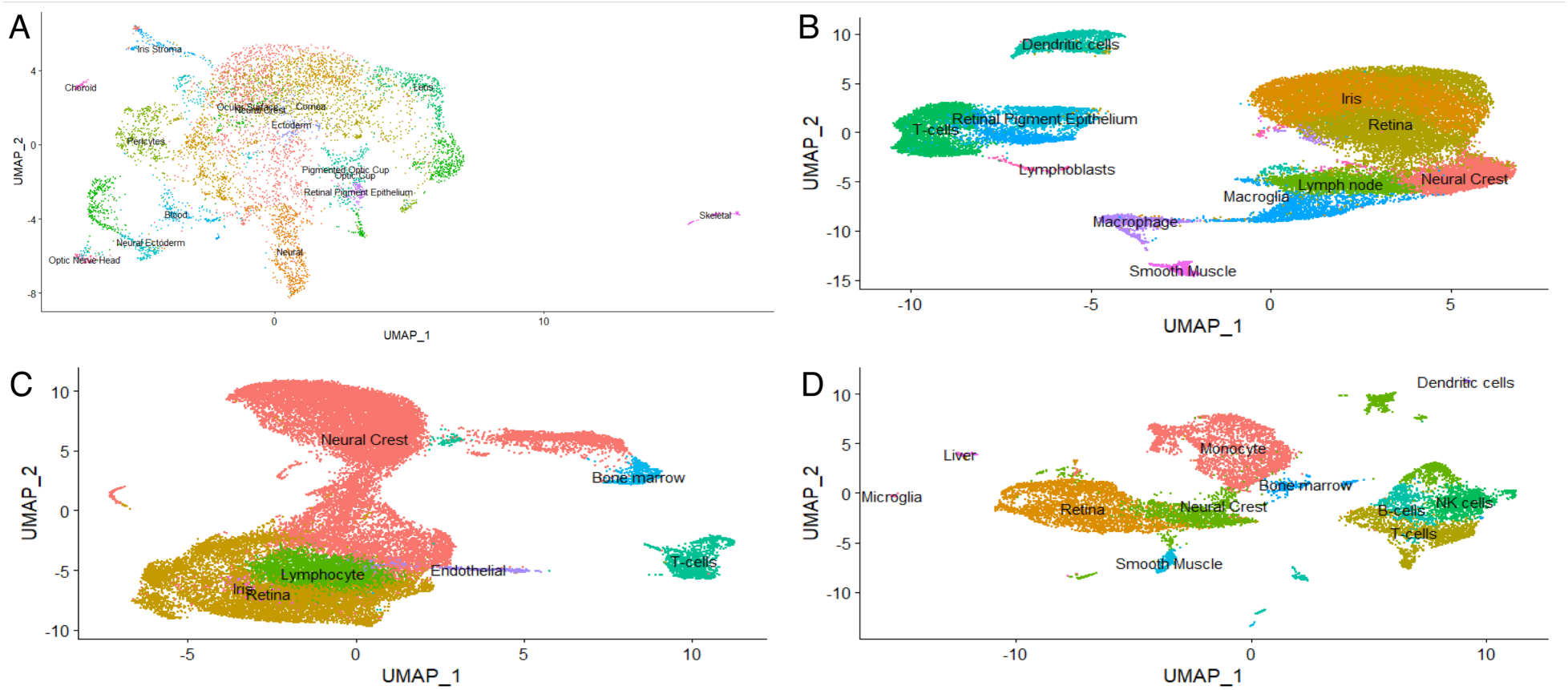
UMAP plots displaying spatial organization. UMAP plots of the (A) SEAM organoid model, (B) UM class 1 metastasis phenotype, (C) UM class 2 primary phenotype, and (D) UM class 2 metastasis phenotype with labeled scRNA-seq analysis clusters. Differentially expressed genes were compared between the SEAM, UM class 1 metastatic, UM class 2 primary, and UM class 2 metastatic clusters.

Representative of human eye tissue, the SEAM organoid should have similar gene expressions, cell type specification, and spatial organization as a normal human eye, adjusting for the fact that most sequenced human eye tissue would be at a more mature eye development stage than the SEAM organoid. Likewise, the scRNA-seq data from UM patients should have gene expression levels, cell type specification, and spatial organization similar to those of the BAP1 KD model. Using the scRNA-seq expression plot data (Figure 4) and differential gene expression plotted through log_2_ fold change, the most expressed genes within the neural crest in the normal human eye, SEAM, and UM samples of various classes were selected, along with other genes with notable functions or significant gene expressions (p<0.05). BAP1 was selected since it is the most commonly mutated tumor suppressor gene in UM class 2 (Masclef, 2021). ABCB5 was selected since it is a neural crest-located UM prognostic factor (Broggi, 2019). STRO1 was selected since it indicates neural crest expression (Ning, 2011). SOX10 was selected as it is a UM marker and transcription factor that mediates downstream neural crest differentiation (Alghamdi, 2015) and is highly expressed in the retina and iris stroma, two of the locations into which neural crest commonly differentiates (Betancur, 2010). KI67 (MKI67) was selected because it indicates general cell proliferation (Li, 2014). AKT1 was selected because it is implicated in UM migration and metastasis (Farhan, 2021). CASP3 was selected because it is a marker for apoptosis (Brentnall, 2013). PRKDC was selected since it is a DNA repair gene that drives angiogenesis and metastatic tumor migration by promoting pro-metastatic protein secretion (Mohiuddin, 2019). p53 (TP53) was selected since it has been shown to aid UM’s invasion ability (Liu, 2017). PAX3 was selected since it is another neural crest marker implicated in aiding UM differentiation and progression (Hathaway, 2011). PGC1α (PPARGC1A) and NRF1 were selected since they are oxidative phosphorylation markers (Patti, 2003). HIF1 was selected since it has been shown to aid UM tumor growth by facilitating angiogenesis under low oxygen conditions (Asnaghi, 2014). SIRT1 was selected since tumors lacking it develop metastasis faster than those with SIRT1 (Oltean, 2009).

**Figure 4:**
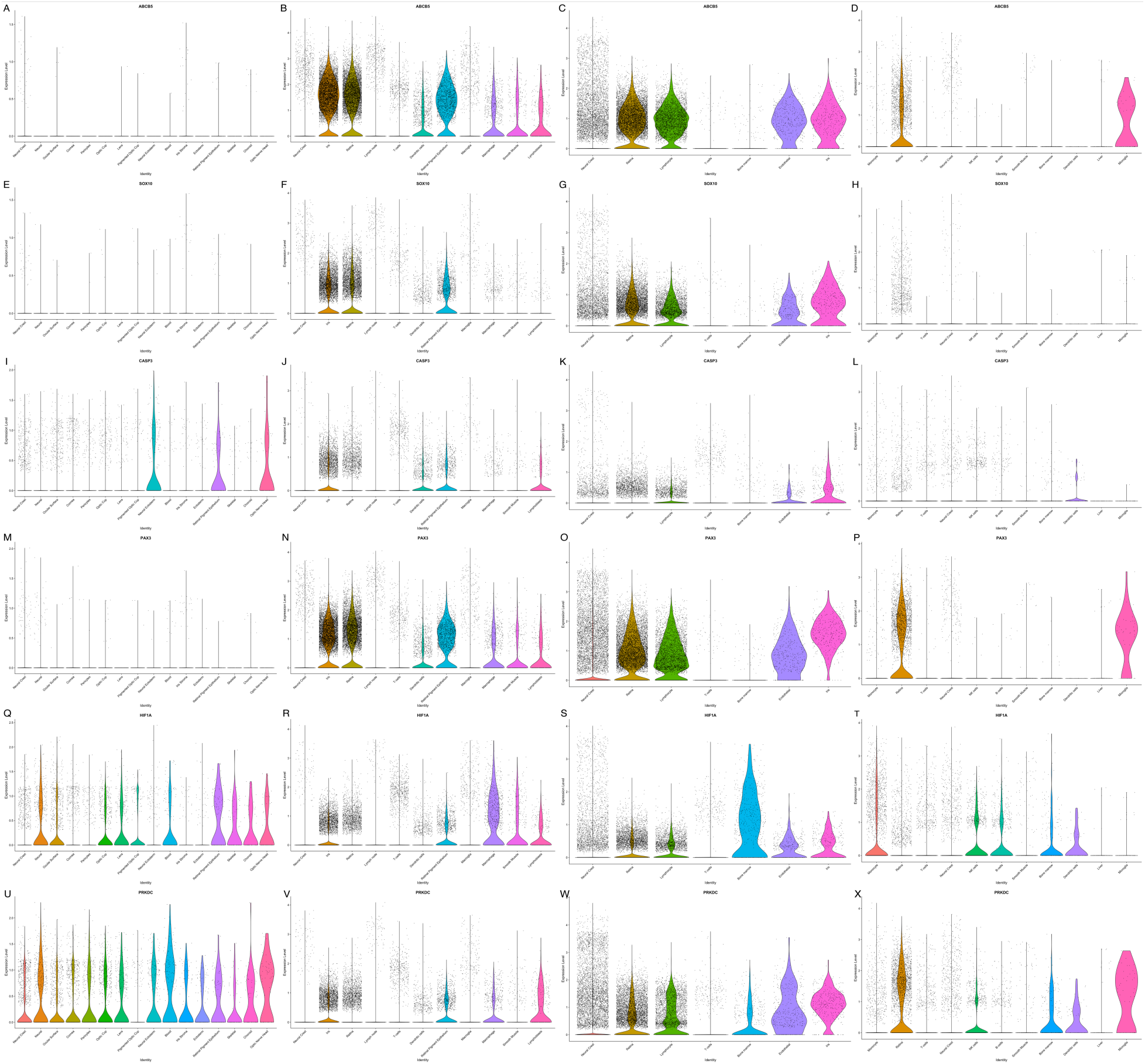
Sample of violin plots depicting significant neural crest gene expression. Clustered gene expressions for (A) SEAM ABCB5, (B) UM class 1 metastatic ABCB5, (C) UM class 2 primary ABCB5, (D) UM class 2 metastatic ABCB5, (E) SEAM SOX10, (F) UM class 1 metastatic SOX10, (G) UM class 2 primary SOX10, (H) UM class 2 metastatic SOX10, (I) SEAM CASP3, (J) UM class 1 metastatic CASP3, (K) UM class 2 primary CASP3, (L) UM class 2 metastatic CASP3, (M) SEAM PAX3, (N) UM class 1 metastatic PAX3, (O) UM class 2 primary PAX3, (P) UM class 2 metastatic PAX3, (Q) SEAM HIF1 (HIF1A), (R) UM class 1 metastatic HIF1 (HIF1A), (S) UM class 2 primary HIF1 (HIF1A), (T) UM class 2 metastatic HIF1 (HIF1A), (U) SEAM PRKDC, (V) UM class 1 metastatic PRKDC, (W) UM class 2 primary PRKDC, and (X) UM class 2 metastatic PRKDC in corresponding SEAM and UM clusters of various classes were compared.

The dCas9-KRAB SEAM organoids (Figure 5) were grown for 35 days until relative maturity. After lentiviral transduction of the CRISPR-Cas9 BAP1 knockdown vectors into some organoids, the control, doxycycline, BAP1 KD, and BAP1 KD + doxycycline cells were stained and analyzed through immunofluorescence (Figure 6) to analyze the BAP1 KD phenotype’s ability to induce uveal melanoma in dCas9 KRAB cells and analyze doxycycline’s effects on a UM phenotype. Immunofluorescence was conducted for proteins indicating total cell count (DAPI), tumor suppression (BAP1), UM prognosis in the neural crest (ABCB5, PAX3, and SOX10), UM metastasis (AKT1), neural crest proliferation (KI67 + STRO1 co-expression), and apoptosis (CASP3). The wells required for each immunofluorescence analysis were grown and repeated five times and then averaged to ensure accurate readings throughout the control vs. doxycycline, control vs BAP1 KD (UM phenotype), control vs BAP1 KD + doxycycline (treated UM phenotype), and BAP1 KD (UM phenotype) vs. BAP1 KD + doxycycline (treated UM phenotype) groups (Figure 7). Lentiviral infection was successful, as indicated by significant BAP1 KD (6G) and GFP in the nucleus (Figures 6K, 6L). For those figures, ABCB5 was indicated by the cell outlines (Figures 6I, 6J, 6K, 6L).

**Figure 5:**
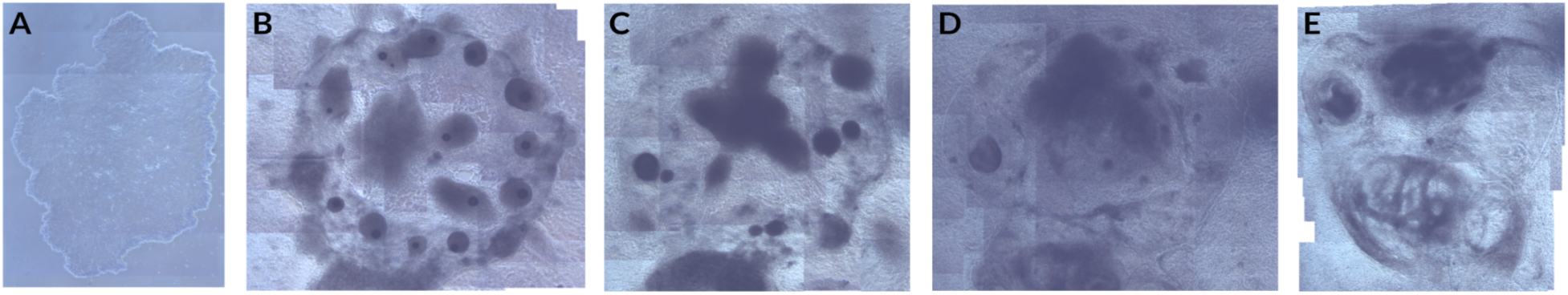
Leica inverted 5x microscope Inkscape (Inkscape Project, 2021) combined photo of the stem cell SEAM culture (passage 67) at (A) 0, (B) 13, (C) 18, (D) 26, and (E) 35 days into development.

**Figure 6:**
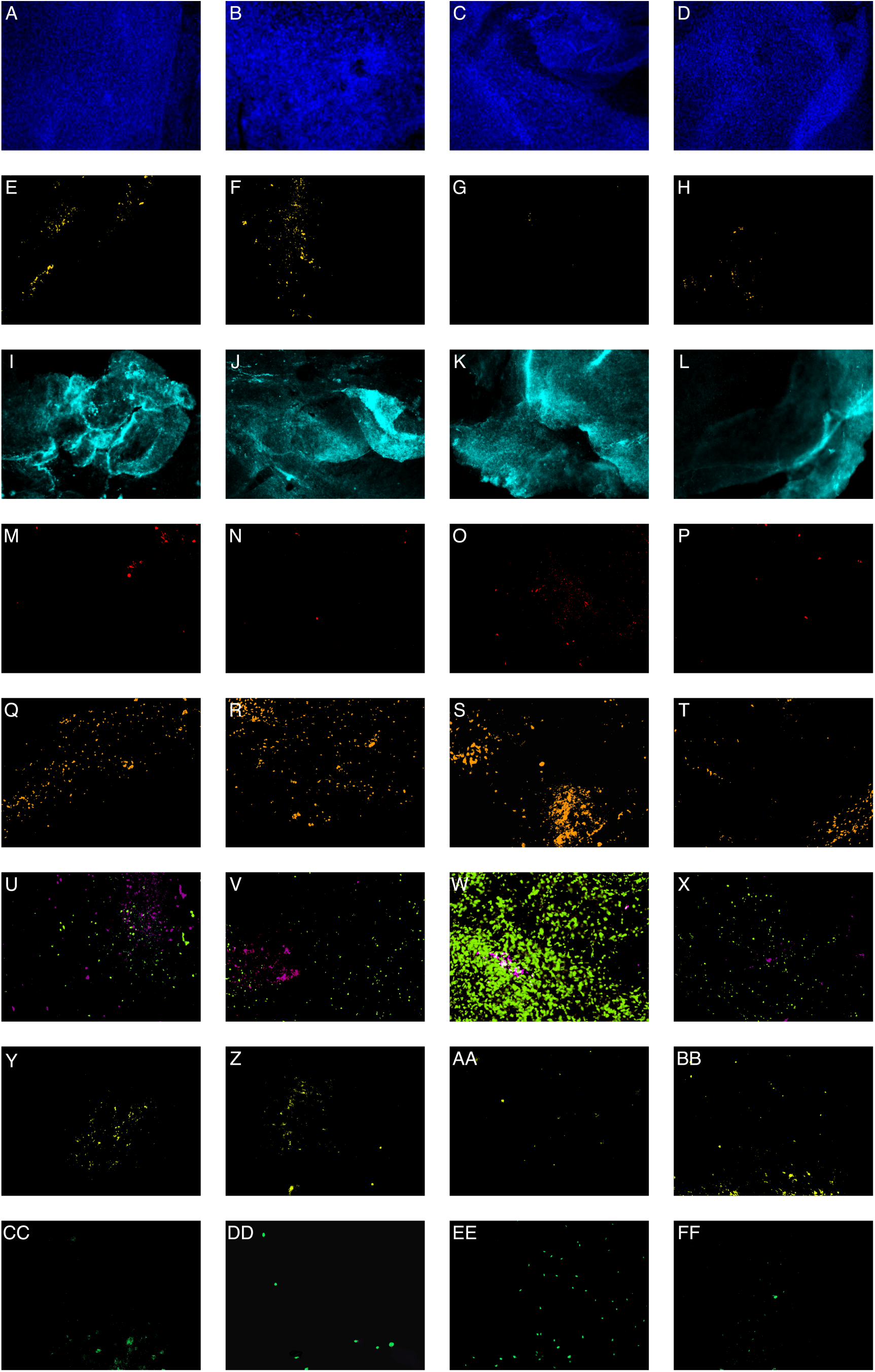
Leica immunofluorescence photos displaying dCas9 KRAB SEAM protein marker expressions after 35 days of development at 50x. Expression of (A) control DAPI, (B) doxycycline DAPI, (C) BAP1 KD DAPI, (D) BAP1 KD + doxycycline DAPI, (E) control BAP1, (F) doxycycline BAP1, (G) BAP1 KD BAP1, (H) BAP1 KD + doxycycline BAP1, (I) control ABCB5 (outlines), (J) doxycycline ABCB5 (outlines), (K) BAP1 KD ABCB5 (outlines), (L) BAP1 KD + doxycycline ABCB5 (outlines), (M) control AKT1, (N) doxycycline AKT1, (O) BAP1 KD AKT1, (P) BAP1 KD + doxycycline AKT1, (Q) control PAX3, (R) doxycycline PAX3, (S) BAP1 KD PAX3, (T) BAP1 KD + doxycycline PAX3, (U) control KI67 + STRO1 co-expression, (V) doxycycline KI67 + STRO1 co-expression, (W) BAP1 KD KI67 + STRO1 co-expression, (X) BAP1 KD + doxycycline KI67 + STRO1 co-expression, (Y) control CASP3, (Z) doxycycline CASP3, (AA) BAP1 KD CASP3, (BB) BAP1 KD + doxycycline CASP3, (CC) control SOX10, (DD) doxycycline SOX10, (EE) BAP1 KD SOX10, and (FF) BAP1 KD + doxycycline SOX10 indicated by colored dots.

**Figure 7:**
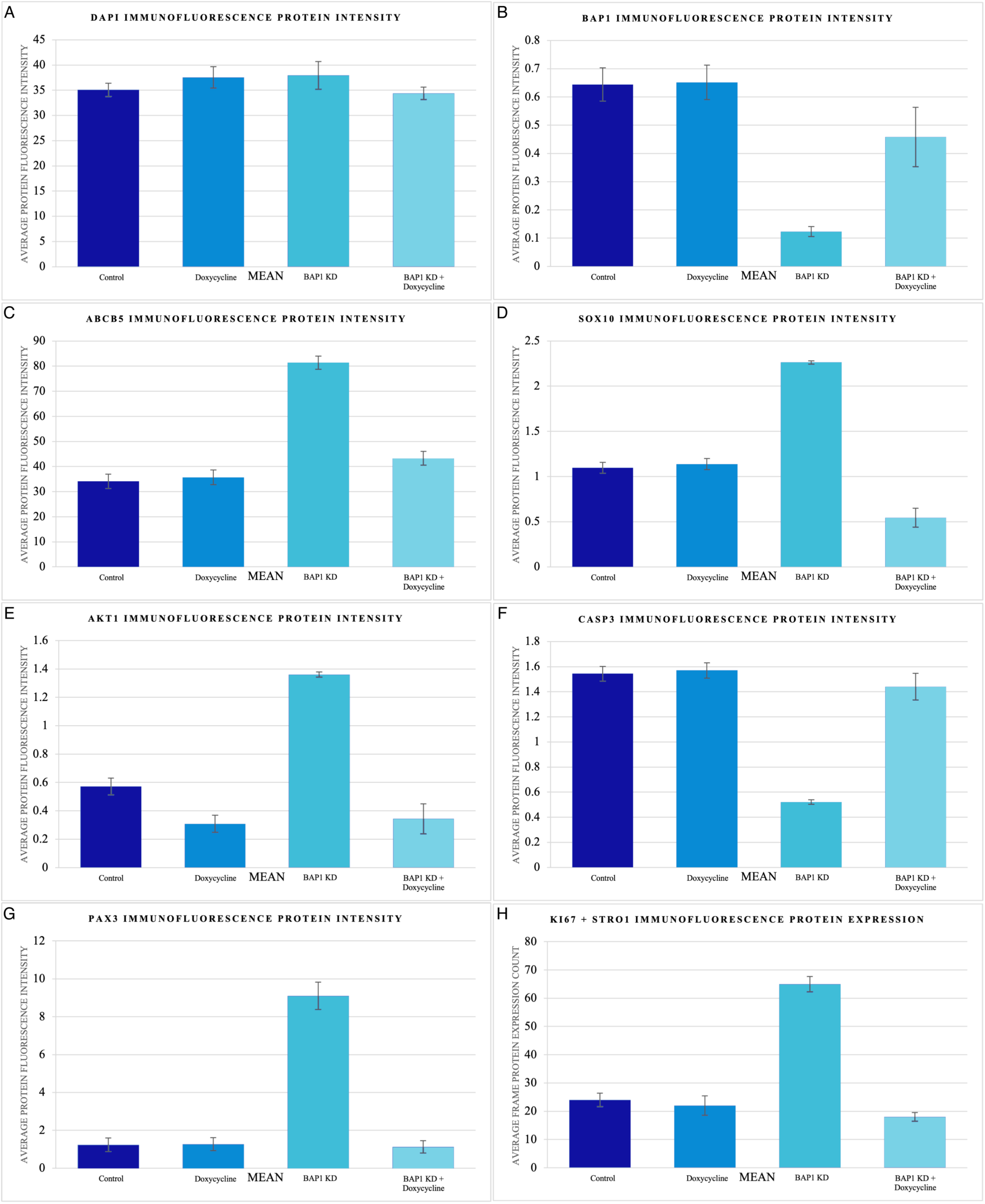
Labeled protein markers in the dCas9 KRAB SEAM organoids analyzed through immunofluorescence and quantified through ImageJ’s average protein fluorescence intensity method. Control, doxycycline, BAP1 KD, and BAP1 KD + doxycycline protein expression levels for (A) DAPI, (B) BAP1, (C) ABCB5, (D) SOX10, (E) AKT1, (F) CASP3, (G) PAX3, and (H) KI67 + STRO1.

In regard to control vs. doxycycline, the doxycycline sample alone exhibited no significant change compared to the control in DAPI, BAP1, ABCB5, PAX3, KI67+STRO1 co-expression, CASP3, and SOX10 protein expression. However, the AKT1 protein was significantly decreased (p<0.05) following doxycycline treatment.

For control vs. BAP1 KD (UM phenotype), the BAP1 KD (UM phenotype) sample exhibited no significant change compared to the control in DAPI expression. However, the BAP1 KD (UM phenotype) sample exhibited significant decreases in BAP1 (p<0.05), increases in ABCB5 (p<0.0001), increases in AKT1 (p<0.001), increases in PAX3 (p<0.001), increases in KI67 + STRO1 co-expression (p<0.05), decreases in CASP3 (p<0.01), and increases in SOX10 (p<0.05) protein expressions.

Regarding control vs. BAP1 KD + doxycycline (treated UM phenotype), the BAP1 KD + doxycycline (treated UM phenotype) sample was not significantly changed for DAPI, BAP1, AKT1, PAX3, KI67 + STRO1 co-expression, and CASP3. Yet compared to the control, SOX10 protein expression was significantly decreased (p<0.05) with BAP1 KD + doxycycline (treated UM phenotype).

As for BAP1 KD (UM phenotype) vs BAP1 KD + doxycycline (treated UM phenotype), the DAPI expression was not significantly changed. Compared to the BAP1 KD (UM phenotype), the BAP1 KD + doxycycline (treated UM phenotype) sample exhibited highly significant increases in BAP1 (p<0.0001), decreases in ABCB5 (p<0.0001), decreases in AKT1 (p<0.001), decreases in PAX3 (p<0.001), decreases in KI67 + STRO1 co-expression (p<0.0001), increases in CASP3 (p<0.01), and decreases in SOX10 (p<0.01) protein expressions.

To confirm the immunofluorescence analysis results and further determine the BAP1 KD phenotype’s ability to induce uveal melanoma in dCas9 KRAB cells and analyze doxycycline’s effects on a UM phenotype, gene primers indicating tumor suppression (BAP1), UM prognosis in the neural crest (ABCB5 and PAX3), angiogenesis (PRKDC and HIF1), UM invasion (p53), oxidative phosphorylation (PGC1α and NRF1), and metastasis inhibition (SIRT1) were quantified using RT-qPCR analysis. RT-qPCR was repeated five times and then averaged to ensure accurate readings throughout the control vs. doxycycline, control vs BAP1 KD (UM phenotype), control vs BAP1 KD + doxycycline (treated UM phenotype), and BAP1 KD (UM phenotype) vs. BAP1 KD + doxycycline (treated UM phenotype) groups (Figure 8).

**Figure 8:**
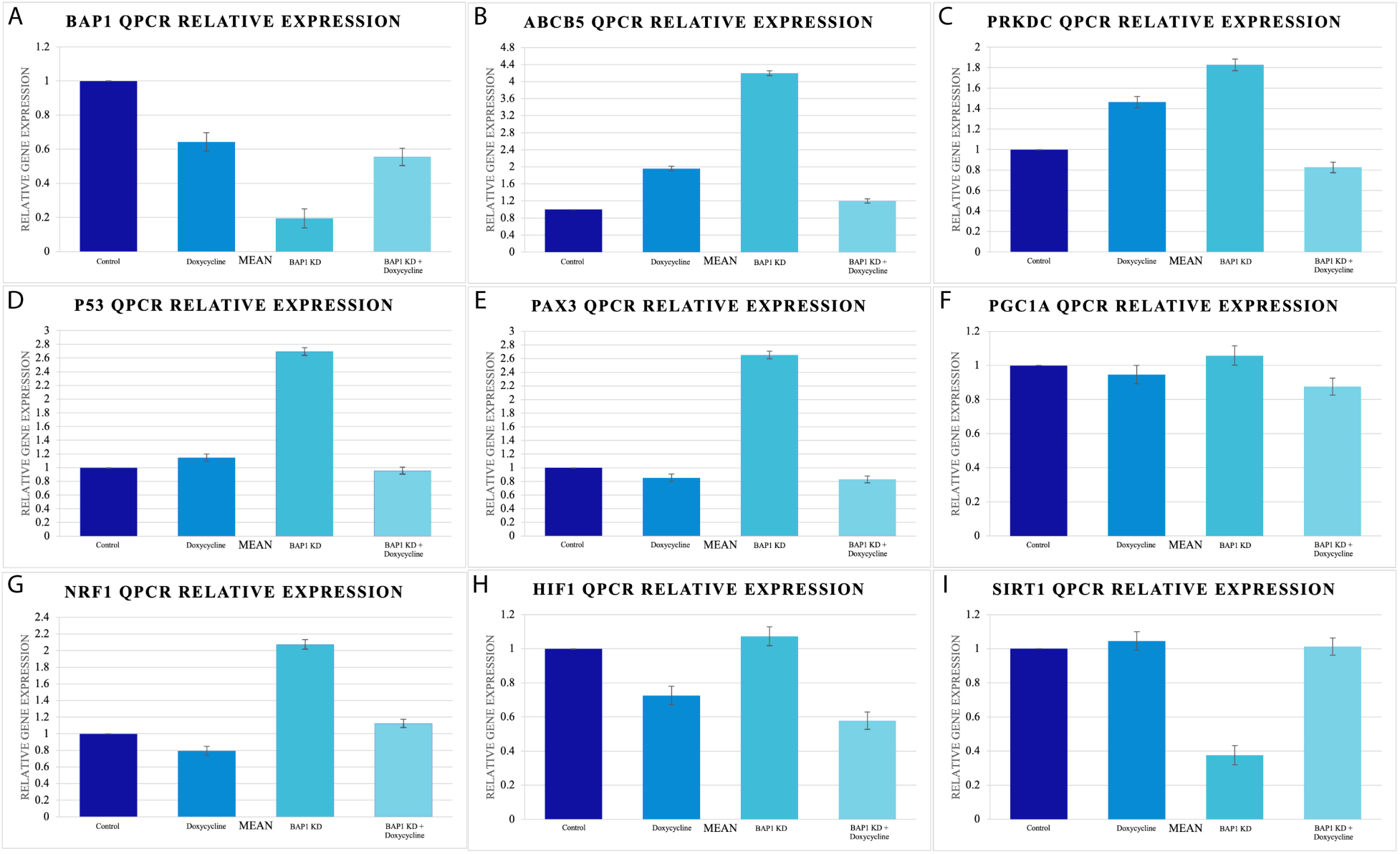
Expression levels for the genes evaluated in the dCas9 KRAB organoids determined based on qPCR cycles and compared to the cyclophilin reference gene for normalization. Control, doxycycline, BAP1 KD, and BAP1 KD + doxycycline gene expression levels for (A) BAP1, (B) ABCB5, (C) PRKDC, (D) p53, (E) PAX3, (F) PGC1α, (G) NRF1, (H) HIF1, and (I) SIRT1.

After doxycycline treatment, the BAP1, ABCB5, PRKDC, p53, PAX3, PGC1α, NRF1, HIF1, and SIRT1 gene expressions analyzed did not exhibit significant deviation compared to the control, indicating minimal side effects for doxycycline treatment without BAP1 KD (UM phenotype).

Following BAP1 KD (UM phenotype), PGC1α and HIF1 gene expressions slightly increased compared to the control, although not by a significant amount. However, the BAP1 KD (UM phenotype) exhibited significant gene expression decreases in BAP1 (p<0.0001), increases in ABCB5 (p<0.05), increases in PRKDC (p<0.05), increases in p53 (p<0.05), increases in PAX3 (p<0.0001), increases in NRF1 (p<0.05), and decreases in SIRT1 (p<0.01) compared to the control.

As for the BAP1 KD + doxycycline (treated UM phenotype), ABCB5, PRKDC, p53, PAX3, NRF1, and SIRT1 did not significantly deviate compared to the control. Yet the BAP1 KD + doxycycline (treated UM phenotype) was significantly decreased for HIF1 (p<0.05) compared to the control.

Regarding the BAP1 KD (UM phenotype) vs BAP1 KD + doxycycline (treated UM phenotype), all gene markers evaluated significantly changed for each condition. Compared to the BAP1 KD (UM phenotype), the BAP1 KD + doxycycline (treated UM phenotype) exhibited significant increases in BAP1 (p<0.05), decreases in ABCB5 (p<0.05), decreases in PRKDC (p<0.05), decreases in p53 (p<0.05), decreases in PAX3 (p<0.0001), decreases in PGC1α (p<0.05), decreases in NRF1 (p<0.05), decreases in HIF1 (p<0.05), and increases in SIRT1 (p<0.05).

## Discussion

The clusters of the SEAM, UM class 1 metastatic, UM class 2 primary, and UM class 2 metastatic were identified through Enrichr, gene function literature reviews, and GeneCards analysis. Thus, the genes corresponding to each cluster were labeled according to their best-fit eye region. Compared to the normal human eye, the SEAM organoid has similar cell type specification and spatial organization among the SEAM clusters, so it accurately represents generalizable neural crest development within the human eye. Consistent with past results that SF3B1 and EIF1AX mutations cause UM class 1 while BAP1 mutations are critical to proliferation in UM class 2 (Durante, 2020), the gene expressions, cell type specification, and spatial organization within the UM clusters evaluated replicated those of typical metastasizing UM tumors.

Thus, the scRNA-seq samples collected from SEAM accurately reflected human eye tissue. Similarly, the scRNA-seq samples collected from the UM patients exhibited comparable gene expressions, cell type specification, and spatial organization to those of the BAP1 KD (UM phenotype) model, further corroborating that the BAP1 KD lentiviral transduction effectively replicates UM based on the BAP1 KD and nucleic GFP data. Additionally, based on the presence of a neural crest cluster, which eventually develops into the regions from which UM metastasizes, the SEAM organoid was further verified as a promising model for analyzing the genetic origin and potential treatments for UM. As a novel human eye replicating model free from the limitations of animal and human cell lines, the SEAM organoid can be used to evaluate treatments for not only UM but also other ophthalmic pathologies. The bioinformatic differences in gene expressions between the control and UM neural crest clusters determined by scRNA-seq (Figure 4), along with all other gene and protein expressions examined in the experiment (BAP1, STRO1, KI67, AKT1, p53, PGC1α, NRF1, and SIRT1), were corroborated *in vitro* through immunofluorescence and RT-qPCR for the control vs. BAP1 KD (UM phenotype) groups. However, although similar significant (p<0.05) genes were identified in the SEAM and UM datasets, which were later analyzed through immunofluorescence and RT-qPCR, targeted therapies like doxycycline seem to exhibit differing effects on those genes for the normal SEAM control organoids vs. BAP1 KD organoids (UM phenotype).

For immunofluorescence and RT-qPCR, the consistent total cell levels (DAPI) in the control and experimental groups control against varying cell levels causing protein expression differences, and all the gene expression levels are normalized against the Cyclophilin A reference gene.

Based on the control vs. doxycycline evaluation utilizing both immunofluorescence and RT-qPCR, doxycycline seems to have insignificant effects compared to the non-UM-phenotype control on total cell count and cytotoxicity (DAPI), tumor suppression (BAP1), UM prognosis in the neural crest (ABCB5, PAX3, and SOX10), neural crest proliferation (KI67+STRO1 co-expression), apoptosis (CASP3), angiogenesis (PRKDC and HIF1), UM invasion (p53), oxidative phosphorylation (PGC1α and NRF1), and metastasis inhibition (SIRT1). Thus, doxycycline likely has minimal cytotoxic effects on normal human eye tissue regarding these factors. However, doxycycline significantly decreased (p<0.05) AKT1 expression, indicating that doxycycline treatment on a control sample may potentially lead to slowed cell proliferation and differentiation.

The control vs. BAP1 KD (UM phenotype) analysis with immunofluorescence and RT-qPCR revealed no significant change in DAPI expression, indicating that total protein expression levels remained relatively constant. With such a baseline, the oxidative phosphorylation (PGC1α) and angiogenesis (HIF1) levels were found to be slightly increased for the BAP1 KD (UM phenotype) compared to the control, although not by a significant amount. While the PGC1α oxidative phosphorylation and HIF1 angiogenesis levels may have slightly increased, albeit insignificantly, they did not decrease, which is notable since both are essential for UM proliferation in the BAP1 KD (UM phenotype) model. After all, oxidative phosphorylation (PGC1α) and angiogenesis (HIF1) aid UM metastasis, regardless of whether their levels stay contract between the control vs. BAP1 KD (UM phenotype) models. However, the BAP1 KD (UM phenotype) exhibited significant increases (p<0.05) in oxidative phosphorylation (NRF1) gene expression, suggesting that the BAP1 KD (UM phenotype) may draw on the NRF1 gene more than the PGC1α gene for oxidative phosphorylation. The BAP1 KD (UM phenotype) also exhibited significant decreases in tumor suppressor (BAP1) protein (p<0.05) and gene (p<0.0001) expression, indicating a successful BAP1 KD creating the UM phenotype. Additionally, the BAP1 KD (UM phenotype) exhibited significant increases in neural crest (ABCB5: p<0.0001, p<0.05; PAX3: p<0.001, p<0.0001; and SOX10: p<0.05, N/A) protein and gene expressions, further indicating a more UM-like phenotype. Furthermore, the BAP1 KD (UM phenotype) had significant increases (p<0.001) in AKT1 protein expression, indicating increased UM metastasis, as AKT1 aids in UM migration. Correspondingly, the BAP1 KD (UM phenotype) exhibited significant increases (p<0.05) in neural crest protein proliferation (KI67 + STRO1 co-expression), indicating increased neural crest-like UM growth. The BAP1 KD (UM phenotype) also exhibited significant decreases (p<0.01) in apoptosis (CASP3) protein expression, indicating inactivated tumor suppressor genes, oncogene activation, and uncontrolled UM proliferation. Additionally, the BAP1 KD (UM phenotype) exhibited significant increases (p<0.05) in angiogenesis (PRKDC) gene expression, likely supporting further UM growth. Furthermore, the BAP1 KD (UM phenotype) had significant increases (p<0.05) in UM invasion (p53) gene expression, corroborating increased UM growth. Simultaneously, the BAP1 KD (UM phenotype) had significant decreases (p<0.01) in metastasis inhibition (SIRT1), indicating unrestricted UM proliferation.

The control vs. BAP1 KD + doxycycline (treated UM phenotype) analysis revealed insignificant protein and gene expression changes for the BAP1 KD + doxycycline (treated UM phenotype) compared to the control regarding total cell count and cytotoxicity (DAPI), tumor suppression (BAP1), UM prognosis in the neural crest (ABCB5 and PAX3), UM metastasis (AKT1), neural crest proliferation (KI67 + STRO1 co-expression), apoptosis (CASP3), angiogenesis (PRKDC), UM invasion (p53), oxidative phosphorylation (PGC1α and NRF1), and SIRT1 expression. However, immunofluorescence revealed that SOX10 neural crest protein expression was significantly decreased (p<0.05) for the BAP1 KD + doxycycline (treated UM phenotype), implying that compared to other neural crest markers like ABCB5, PAX3, and STRO1, doxycycline may target SOX10 more when exposed to the BAP1 KD (UM phenotype) model. Such a decrease in SOX10 phenomenon may potentially affect post-UM MITF activation and, subsequently, melanocyte production. Additionally, SOX10 is a marker for stem cell development, although its role in mediating stem cell development may not be as relevant for the human eye once the eye has fully matured, as is the case in most uveal melanoma cases. Furthermore, RT-qPCR revealed that the HIF1 angiogenesis gene marker expression was also significantly decreased (p<0.05) for the BAP1 KD + doxycycline (treated UM phenotype), contrary to the insignificant change indicated by the PRKDC angiogenesis marker. Doxycycline-treated BAP1 KD (UM phenotype) plausibly targets HIF1 more than PRKDC in angiogenesis, causing HIF1 expression to be significantly decreased (p<0.05) compared to PRKDC. Such a decrease in HIF1 may potentially affect post-UM-treatment metabolic oxygen homeostasis during hypoxia, possibly worsening post-UM-treatment low oxygen tolerability.

As for BAP1 KD (UM phenotype) vs. BAP1 KD + doxycycline (treated UM phenotype), the total cell count (DAPI) protein levels did not significantly change, indicating relatively constant total cell expression. Thus, it is unlikely that doxycycline had cytotoxic effects leading to decreased cell counts. Compared to the BAP1 KD (UM phenotype), the BAP1 KD + doxycycline (treated UM phenotype) protein and gene expressions exhibited significant increases in tumor suppression (BAP1: p<0.0001, p<0.05) and decreases in UM prognosis in the neural crest (ABCB5: p<0.0001, p<0.05; PAX3: p<0.001, p<0.0001; and SOX10: p<0.01, N/A). Additionally, the BAP1 KD + doxycycline (treated UM phenotype) protein and gene expressions had significant decreases (p<0.0001) in neural crest proliferation (KI67 + STRO1), decreases (p<0.001) in UM metastasis (AKT1), increases (p<0.01) in apoptosis (CASP3), decreases (p<0.05, p<0.05) in angiogenesis (PRKDC and HIF1), decreases (p<0.05) in UM invasion (p53), decreases (p<0.05, p<0.05) in oxidative phosphorylation (PGC1α and NRF1), and increases (p<0.05) in metastasis inhibition (SIRT1). The significant decrease in PGC1α for the BAP1 KD + doxycycline (treated UM phenotype) compared to the BAP1 KD (UM phenotype) but not for the other comparative phenotypes may plausibly be caused by doxycycline targeting oxidative phosphorylation areas modulated by the NRF1 gene more than those of the PGC1α gene. Decreased NRF1 expression may limit oxidative phosphorylation, thereby significantly decreasing PGC1α expression, which would cause PGC1α to significantly decrease even if doxycycline did not directly affect PGC1α. This phenomenon is notable since PGC1α does not significantly increase for the BAP1 KD (UM phenotype) compared to the control, but NRF1 does.

Considering doxycycline’s inhibitory effects on oxidative phosphorylation, angiogenesis, neural crest proliferation, and UM growth for the UM phenotype, doxycycline seems to be a viable candidate for UM treatment. Uniquely, doxycycline seems to have more drastic effects on the BAP 1 KD (UM phenotype) organoids rather than the control normal human eye organoids, perhaps due to doxycycline targeting proliferating tumors that require more mitochondrial oxidative phosphorylation due to their higher energy consumption. Corroborating that hypothesis, the more significant decreases in angiogenesis and UM proliferation genes compared to the insignificant decrease caused by doxycycline in control normal human eye organoids indicates that doxycycline does not have as drastic effects in the control human eye organoids as it does in the BAP1 KD (UM phenotype) organoids. However, given that doxycycline significantly decreases AKT1 expression in the control, potentially leading to slowed cell proliferation and differentiation, doxycycline should not be used as a preventative treatment for UM since it may negatively affect those with pre-existing slow cell growth conditions.

Since the BAP1 KD + doxycycline (treated UM phenotype) group did not significantly affect most protein and gene expression levels, doxycycline treatment likely restored a relatively normal phenotype following BAP1 KD (UM phenotype). However, the BAP1 KD + doxycycline (treated UM phenotype) exhibited significant decreases in SOX10, potentially inhibiting melanocyte production, and HIF1, possibly worsening hypoxia symptoms. While the scope of this study focused on doxycycline’s short-term effects over 14 days, doxycycline’s long-term effects on the BAP1 KD (UM phenotype) are yet undetermined, along with doxycycline’s effects over less than 14 days. Regardless, after the tumor-suppressing effects have materialized, the BAP1 KD + doxycycline (treated UM phenotype) will likely need to wean off the doxycycline doses provided every 48 hours to restore normal cell function.

## Conclusions

As the SEAM organoid model resembles the human eye by scRNA-seq (gene expressions, cell type specification, and spatial organization) and immunofluorescence, further treatments for UM or other ophthalmic pathologies can also be evaluated free from the limitations of animal and human cell lines. However, to ensure complete SEAM organoid maturation, an oxygenated and nutrient-conducive vasculature for the organoid will be further developed through perfusing spinach leaves (Gershlak, 2017), facilitating the continued growth and enlargement of the organoid past 35 days.

The BAP1 KD (UM phenotype) model was found to induce UM metastasis effectively, as 14 days following BAP1 KD, notable markers for oxidative phosphorylation, angiogenesis, neural crest proliferation, and UM growth significantly increased (p<0.05) compared to the control based on the immunofluorescence and RT-qPCR analysis. Gene expression, cell type specification, and spatial organization determined through scRNA-seq analysis corroborated that the BAP1 KD (UM phenotype) model accurately modeled human UM proliferation. Thus, the novel BAP1 KD (UM phenotype) model created in this study effectively replicated UM growth *in vitro,* and has substantial potential as a preliminary model from which the effects of various potential therapies can be evaluated on UM.

After evaluating and comparing SEAM and UM tissue samples through scRNA-seq, unique differentially expressed genes with notable functions were identified. Doxycycline’s effect on those genes was then evaluated through immunofluorescence and RT-qPCR. Utilizing a control SEAM organoid treated with doxycycline, immunofluorescence, and RT-qPCR analyses indicated that the protein expression for the UM metastasis marker AKT1 significantly decreased (p<0.05). In contrast, other oxidative phosphorylation, angiogenesis, neural crest proliferation, and UM growth markers did not significantly change. While doxycycline significantly downregulated the expression of AKT1, which has been implicated in UM metastasis, such downregulation may potentially cause harm by slowing cell proliferation and differentiation. This indicates that doxycycline should only be used after detecting UM growth, not as a preventative therapy. However, aside from AKT1, doxycycline seems to have minimal cytotoxic side effects and can likely be used to treat UM effectively. Notably, the BAP1 KD + doxycycline (treated UM phenotype) exhibited significant decreases in UM prognosis in the neural crest (ABCB5, PAX3, and SOX10), neural crest proliferation (KI67 + STRO1), angiogenesis (PRKDC and HIF1), UM invasion (p53), oxidative phosphorylation (PGC1α and NRF1), along with significant increases in tumor suppression (BAP1), apoptosis (CASP3), and metastasis inhibition (SIRT1). The UM metastasis marker AKT1 also decreased significantly in the BAP1 KD (UM phenotype) vs. BAP1 KD + doxycycline (treated UM phenotype) models, akin to the significant AKT1 decreases exhibited in the control vs. doxycycline models. Consequently, doxycycline seems to target the proliferating UM cells that require more energy consumption and correspondingly more mitochondrial oxidative phosphorylation. To confirm these findings, further expression analyses will be conducted on notable gene and protein markers, and mitochondrial respiration along with glycolysis levels will be evaluated with an Agilent Seahorse XF Analyzer.

Following doxycycline’s application to the BAP1 KD (UM phenotype) model, it was found to mostly restore UM to a relatively normal phenotype. While the doxycycline treatment was successful overall in inhibiting UM, the BAP1 + doxycycline (treated UM phenotype) model still had significantly decreased SOX10, an indicator of UM prognosis in the neural crest, and HIF1, an angiogenesis marker, compared to the control. Thus, melanocyte development may be inhibited by SOX10 downregulation, and hypoxia symptoms may be worsened by HIF1 downregulation.

This study evaluated the short-term effects of BAP1 KD + doxycycline (treated UM phenotype) over 14 days, and shorter- or longer-term studies may be needed. Perhaps more prolonged exposure to BAP1 KD (UM phenotype) lentiviral vectors (over 14 days) would further increase oxidative phosphorylation (PGC1α) and angiogenesis (HIF1) levels compared to the control, and perhaps shorter exposure would alleviate BAP1 KD + doxycycline (treated UM phenotype)’s side effects. Future studies will determine the optimal duration needed for doxycycline-treated BAP1+p53 KD SEAM to return to normal mitochondrial function after treatment by evaluating RT-qPCR oxidative phosphorylation markers and Agilent Seahorse XF Analyzer results. Overall, the trajectories of the doxycycline, BAP1 KD (UM phenotype), and BAP1 KD + doxycycline (treated UM phenotype) may change short- or long-term, alleviating or worsening trends evident in the 14-day short-term study period.

The experiment may also be repeated in the future to study doxycycline’s effects on more notable markers identified with the scRNA-seq analysis. Varying amounts of doxycycline application and differing durations of maturation periods will also be tested for the control and BAP1 KD (UM phenotype) organoids to determine the optimal conditions under which doxycycline is successful. Moreover, since 95% of UM metastasis is to the liver (Bellerive, 2018), doxycycline’s effects will also be evaluated downstream in hepatic cancer. Eventually, the goal is to improve UM outcomes through clinical trials on human patients and alleviate the suffering that UM brings to patients and their families.

Considering that metastatic UM has a current mortality rate of 80% with a six-month median survival period, and currently available UM therapies are both expensive and ineffective, the prescription antibiotic doxycycline seems to be a prime therapeutic candidate for UM that is effective, efficient, and economical.

## Data Availability

Published UM scRNA-seq data were obtained from dbGaP study phs001861.v1.p1 and the associated Durante et al. (2020) study. Complete accessions for the H9 SEAM and normal-eye reference datasets, raw immunofluorescence files, replicate-level quantification, raw RT-qPCR files, and analysis code are available upon request.

## Research Materials and Ethics

This study used established human pluripotent stem cell lines and previously generated deidentified or controlled-access sequencing datasets. No participants were prospectively enrolled for the work reported here.

## Acknowledgements

The author extends special gratitude to Dr. Timothy Blenkinsop for his guidance throughout the author’s scientific journey, Dr. Anne Zebitz-Eriksen, who provided the scRNA-seq datasets used in the report and introduced the author to cell culture and immunostaining techniques, Dr. Rachael Warrington for her advice in various laboratory procedures, and Dr. Bar Makovoz for her guidance in scRNA-seq methods and techniques.

## Appendix

**Supplemental Table 1:**
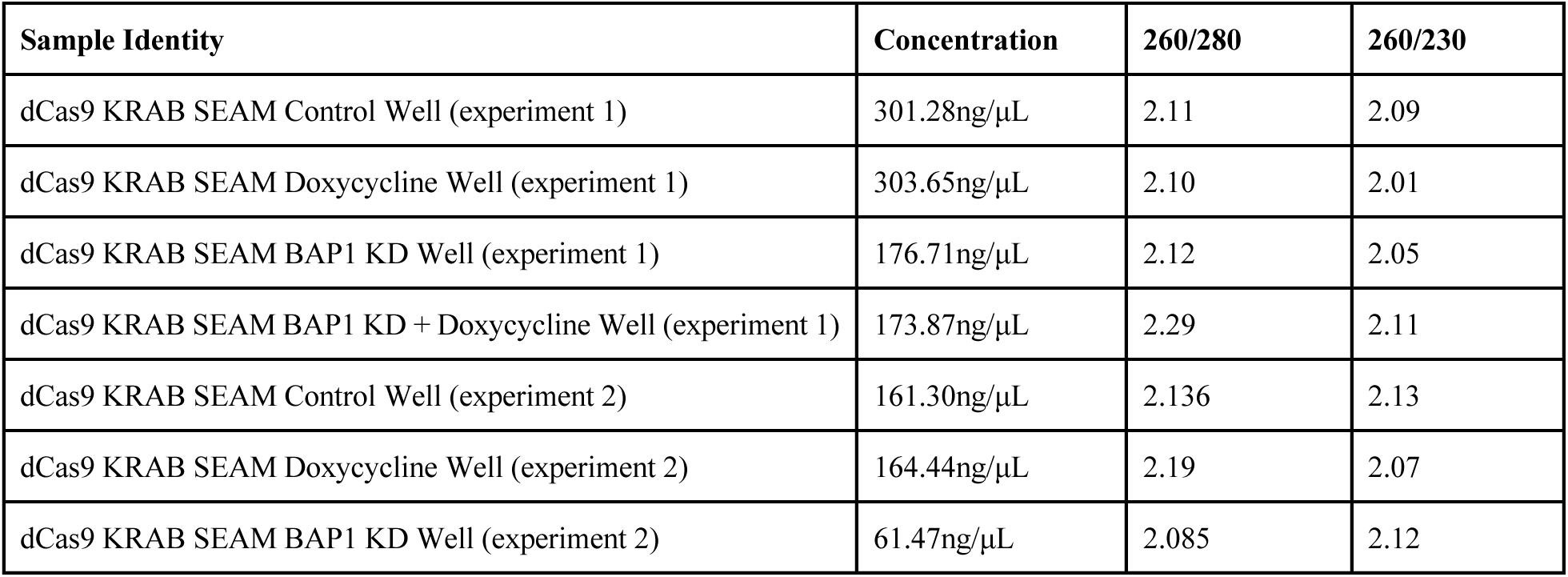
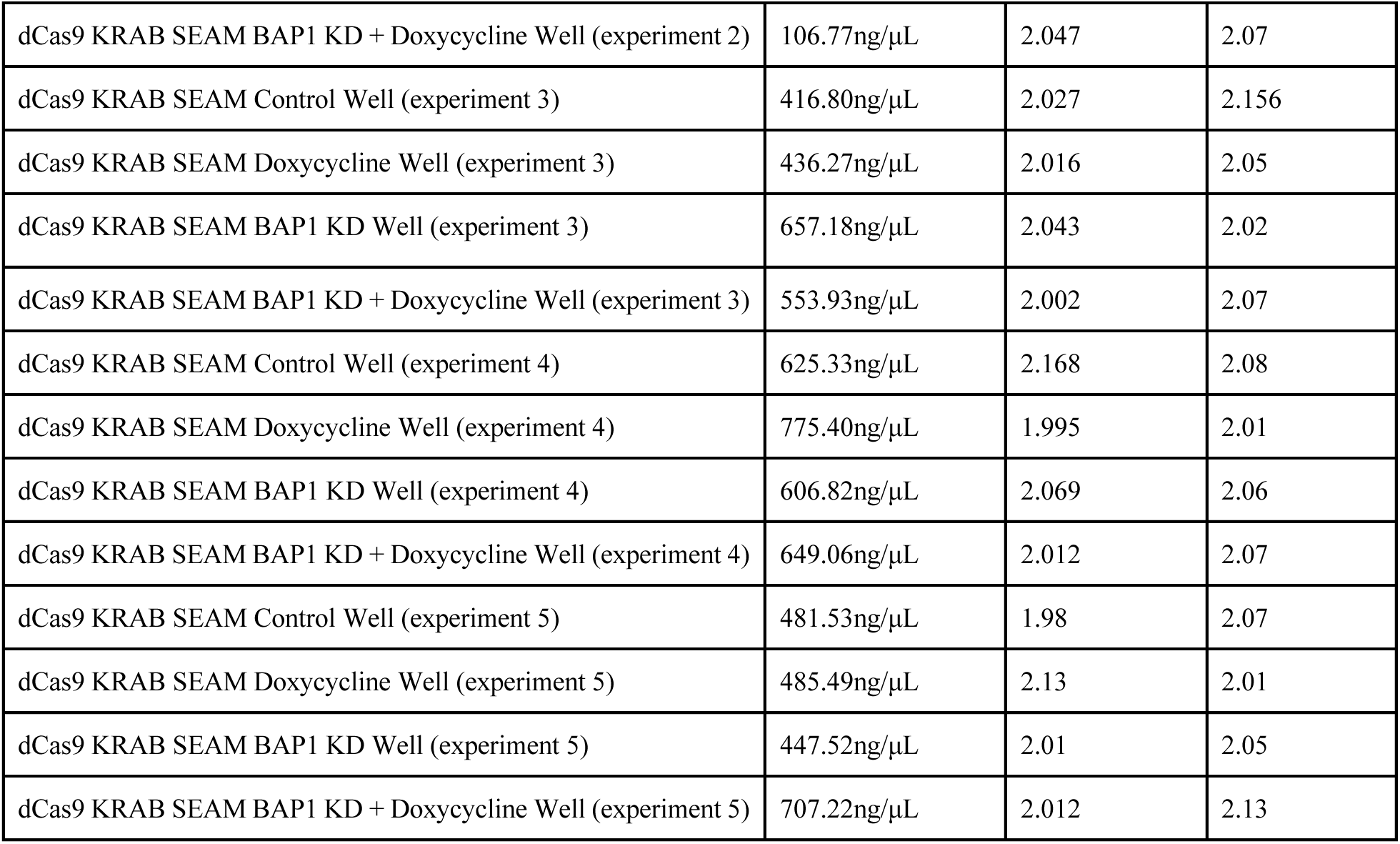
RNA concentration values from NanoDrop ND-1000 spectrophotometer. This spectrophotometry test was run to examine the concentration and purity of RNA in each sample. The 260/280 ratio indicates contaminating proteins, with a pure sample having a value of 2.00. The 260/230 ratio indicates contaminating organic compounds, with a pure sample having a value over 2 (Matlock, 2015).

| Sample Identity | Concentration | 260/280 | 260/230 |
| --- | --- | --- | --- |
| dCas9 KRAB SEAM Control Well (experiment 1) | 301.28ng/μL | 2.11 | 2.09 |
| dCas9 KRAB SEAM Doxycycline Well (experiment 1) | 303.65ng/μL | 2.10 | 2.01 |
| dCas9 KRAB SEAM BAP1 KD Well (experiment 1) | 176.71ng/μL | 2.12 | 2.05 |
| dCas9 KRAB SEAM BAP1 KD + Doxycycline Well (experiment 1) | 173.87ng/μL | 2.29 | 2.11 |
| dCas9 KRAB SEAM Control Well (experiment 2) | 161.30ng/μL | 2.136 | 2.13 |
| dCas9 KRAB SEAM Doxycycline Well (experiment 2) | 164.44ng/μL | 2.19 | 2.07 |
| dCas9 KRAB SEAM BAP1 KD Well (experiment 2) | 61.47ng/μL | 2.085 | 2.12 |
| dCas9 KRAB SEAM BAP1 KD + Doxycycline Well (experiment 2) | 106.77ng/μL | 2.047 | 2.07 |
| dCas9 KRAB SEAM Control Well (experiment 3) | 416.80ng/μL | 2.027 | 2.156 |
| dCas9 KRAB SEAM Doxycycline Well (experiment 3) | 436.27ng/μL | 2.016 | 2.05 |
| dCas9 KRAB SEAM BAP1 KD Well (experiment 3) | 657.18ng/μL | 2.043 | 2.02 |
| dCas9 KRAB SEAM BAP1 KD + Doxycycline Well (experiment 3) | 553.93ng/μL | 2.002 | 2.07 |
| dCas9 KRAB SEAM Control Well (experiment 4) | 625.33ng/μL | 2.168 | 2.08 |
| dCas9 KRAB SEAM Doxycycline Well (experiment 4) | 775.40ng/μL | 1.995 | 2.01 |
| dCas9 KRAB SEAM BAP1 KD Well (experiment 4) | 606.82ng/μL | 2.069 | 2.06 |
| dCas9 KRAB SEAM BAP1 KD + Doxycycline Well (experiment 4) | 649.06ng/μL | 2.012 | 2.07 |
| dCas9 KRAB SEAM Control Well (experiment 5) | 481.53ng/μL | 1.98 | 2.07 |
| dCas9 KRAB SEAM Doxycycline Well (experiment 5) | 485.49ng/μL | 2.13 | 2.01 |
| dCas9 KRAB SEAM BAP1 KD Well (experiment 5) | 447.52ng/μL | 2.01 | 2.05 |
| dCas9 KRAB SEAM BAP1 KD + Doxycycline Well (experiment 5) | 707.22ng/μL | 2.012 | 2.13 |

